# Simple Design for Membrane-Free Microphysiological Systems to Model the Blood-Tissue Barriers

**DOI:** 10.1101/2023.10.20.563328

**Authors:** Ashlyn T. Young, Halston Deal, Gabrielle Rusch, Vladimir A. Pozdin, Ashley C. Brown, Michael Daniele

**Affiliations:** Joint Department of Biomedical Engineering, North Carolina State University and University of North Carolina, Chapel Hill, 911 Oval Dr., Raleigh NC, 27695 (USA); Comparative Medicine Institute, North Carolina State University, 1060 William Moore Dr., Raleigh, NC 27606, USA; Department of Electrical & Computer Engineering, Florida International University, Miami, FL (USA); Department of Mechanical & Materials Engineering, Florida International University, Miami, FL (USA); Department of Electrical & Computer Engineering, North Carolina State University, 890 Oval Dr., Raleigh NC, 27695 (USA)

**Keywords:** Microphysiological system, tissue barrier, blood-brain barrier, endothelial cells, microfluidics

## Abstract

Microphysiological systems (MPS) incorporate physiologically relevant microanatomy, mechanics, and cells to mimic tissue function. Reproducible and standardized *in vitro* models of tissue barriers, such as the blood-tissue interface (BTI), are critical for next-generation MPS applications in research and industry. Many models of the BTI are limited by the need for semipermeable membranes, use of homogenous cell populations, or 2D culture. These factors limit the relevant endothelial-epithelial contact and 3D transport, which would best mimic the BTI. Current models are also difficult to assemble, requiring precise alignment and layering of components. The work reported herein details the engineering of a BTI-on-a-chip (BTI Chip) that addresses current disadvantages by demonstrating a single layer, membrane-free design. Laminar flow profiles, photocurable hydrogel scaffolds, and human cell lines were used to construct a BTI Chip that juxtaposes an endothelium in direct contact with a 3D engineered tissue. A biomaterial composite, gelatin methacryloyl and 8-arm polyethylene glycol thiol, was used for *in situ* fabrication of a tissue structure within a Y-shaped microfluidic device. To produce the BTI, a laminar flow profile was achieved by flowing a photocurable precursor solution alongside phosphate buffered saline. Immediately after stopping flow, the scaffold underwent polymerization through a rapid exposure to UV light (<300 mJ·cm^-2^). After scaffold formation, blood vessel endothelial cells were introduced and allowed to adhere directly to the 3D tissue scaffold, without barriers or phase guides. Fabrication of the BTI Chip was demonstrated in both an epithelial tissue model and blood-brain barrier (BBB) model. In the epithelial model, scaffolds were seeded with human dermal fibroblasts. For the BBB models, scaffolds were seeded with the immortalized glial cell line, SVGP12. The BTI Chip microanatomy was analyzed *post facto* by immunohistochemistry, showing the uniform production of a patent endothelium juxtaposed with a 3D engineered tissue. Fluorescent tracer molecules were used to characterize the permeability of the BTI Chip. The BTI Chips were challenged with an efflux pump inhibitor, cyclosporine A, to assess physiological function and endothelial cell activation. Operation of physiologically relevant BTI Chips and a novel means for high-throughput MPS generation was demonstrated, enabling future development for drug candidate screening and fundamental biological investigations.

**HIGHLIGHTS:**

- Barrier-type organs-on-a-chip are popular due to their mimicry of a variety of tissue constructs and interfaces.
- Typical barrier-type organs-on-a-chip rely upon microperforated membranes and complex assembly, which limits both ease of fabrication the desired barrier performance.
- A membrane-free barrier-type organ-on-a-chip is designed, which uses simple Y-channel microfluidics and photopolymerization to form a precise “blood-tissue interface.”
- Fabrication of the membrane-free design can be easily parallelized and scaled-up.

## 1. INTRODUCTION

Microphysiological systems (MPS) are a group of technologies that aim to mimic the structure and function of organs and organ systems to simulate human physiology better than traditional *in vitro* cell cultures. MPS are typically microfluidic devices that contain living cells, extracellular matrix (ECM) or synthetic scaffolds, other biological components, and external stimuli, *e*.*g*., fluid flow, hydrostatic pressure, or electrical stimulation, that mimic the complexity of the natural tissue or organ. An increasing number of MPS designs are being developed as competitive alternatives for fundamental biological investigations and advanced drug development efforts [1]. The field of microfluidic systems and microchannel fabrication has seen significant progress in the past few decades thanks to a multitude of notable researchers [2-4]. MPS to model both developing and pathological lung, kidney, liver, intestine, brain, placenta, skin, and others have been explored. [5-10] Current state of the art MPS designs aim to replicate not only human organs but systemic level “human on a chip” models.[11, 12] Recent advancements have also led to MPS-based products emerging on the commercial market [13].

Among the various MPS designs, accurate models of tissue barriers are particularly crucial to accelerating drug discovery and screening, disease modelling, and pathogenesis. MPS of tissue barriers mimic the function of natural microanatomies, such as the blood-brain barrier [14, 15], skin [16], or intestinal epithelium [17, 18]. The blood-tissue interface, which constitutes a complex barrier separating the blood and tissue compartments, plays a pivotal role in drug discovery and development by enabling the study of systemic transport and signaling throughout the body. High-throughput screening of transport kinetics or barrier interactions can offer significant predictive power for candidate efficacy, prior to animal or human trials. MPS can be utilized to investigate the transport of drugs across individual barriers, providing insights into how treatments can be tailored to individual patients based on their specific barrier characteristics. This approach can be especially beneficial for understanding how drugs are metabolized by different patients and how patient-specific physiology and pharmacokinetics can influence drug efficacy. Compared to traditional in vitro cell culture or in vivo animal models, MPS offer a more accurate and relevant model of the human body. For example, the highly selective blood-brain barrier (BBB) limits the transport of drugs from the bloodstream to the central nervous system, posing a significant challenge to drug development. Therefore, significant effort has been invested in developing MPS to model the BBB. Despite these efforts, many drug candidates still fail to cross the BBB [19]. Moreover, MPS can aid in the evaluation of the toxicity of chemicals and other compounds that may interact with the barrier, which adds value to the safety assessment of consumer products, as well as in the development of new drugs and other chemical compounds.

MPS of the blood-tissue interface typically mimic conventional Transwell® devices and consist of a layered structure of cells and extracellular matrix (ECM) materials [20-22]. They are used to study the transport of molecules across the barrier and the interactions between the barrier and its surrounding environment. Tissue-level function is controlled by transport mechanisms regulated by the direct (contact-dependent) and indirect (paracrine and endocrine) signaling between cells. Thus far, MPS have achieved a high degree of physiological relevance and informative capacity by supporting communication in coculture, even when cell types are separated by a porous, synthetic barrier. To date, some MPS that mimic barrier properties are hand-crafted, multi-layer devices that do not lend themselves to rapid and scalable production or operation. Although there are some industrially produced MPS such as from Mimetas or Emulate, Inc., there is more to be explored in design iterations of new MPS for enhanced barrier simulation [11-13]. The synthetic barrier requirement arises from the challenge to fabricate the microscale structures that are required to mimic the structure and function of the tissue barrier. This can include developing appropriate materials and techniques for creating the layered structure of the barrier, as well as incorporating the necessary microscale channels and sensors for measuring transport and other characteristics of the barrier. However, as this technology matures, it is expected that more MPS will be adopted and on a larger scale in the pharmaceutical industry. Achieving this will be a challenge as the technology must be adapted to meet industrial requirements in terms of cost and scalability.

Innovative fabrication techniques have evolved for MPS designs, including the use of photoactivatable hydrogels based on polyethylene glycol (PEG) [2]. These hydrogels can be patterned and crosslinked using light activation, enabling precise control over the formation of microchannels [23]. This approach offers the unique advantage of greater flexibility and tunability in microchannel design. By manipulating light exposure, researchers can precisely define the geometry and connectivity of the microchannels within the hydrogel. Certain MPS are designed to mimic intricate vascular networks such as the “Y-shaped” channel [24]. This channel configuration offers a more sophisticated model for studying fluid flow dynamics and cellular behavior in a controlled environment. The integration of Y-shaped channels into microfluidic systems provides a valuable tool for tissue engineering and drug testing and adds value to the development of organ-on-a-chip models. The combined benefits of photoactivatable hydrogels and Y-shaped channels motivated the MPS construction methods described in our research.

Herein, we present an MPS that exploits Y-channel microfluidics and stop-flow photolithography to produce a BTI Chip by the hydrodynamic juxtaposition of blood and tissue compartments followed by *in situ* polymerization of a 3D scaffold. This design supports the formation of an endothelium in direct interface with a desired cell population of the tissue compartment while eliminating synthetic barriers or phase guide features, such as micropillars. This dramatically simplifies the fabrication process and tool requirements. The Y-shape of the channel allows alignment of opposing tissue and endothelial sections, which enables the ability to mimic the transport of drugs across the barrier. The simple microfluidic design eliminates the need for multi-step assembly, alignment, or synthetic barriers. The size of the microfluidic channels and relative compartment dimensions provided for the use of 3D printed molds. This design, as shown in **Figure 1**, will eliminate the need for photolithography tools, making device manufacturing more accessible. As previously demonstrated, features in the Y-Channel can be easily scaled by controlling the fluid flow rates [25]. These design features can improve accessibility and manufacturability throughout the models’ adoption by researchers. Upon validating this fabrication technique, we demonstrated the *in situ* formation and characterized the small molecule permeability and regulation of an endothelial-epithelial barrier and blood-brain barrier model. Simultaneous fabrication of multiple MPS was achieved as a proof-of-concept for future scale-up.

**Fig. 1.**
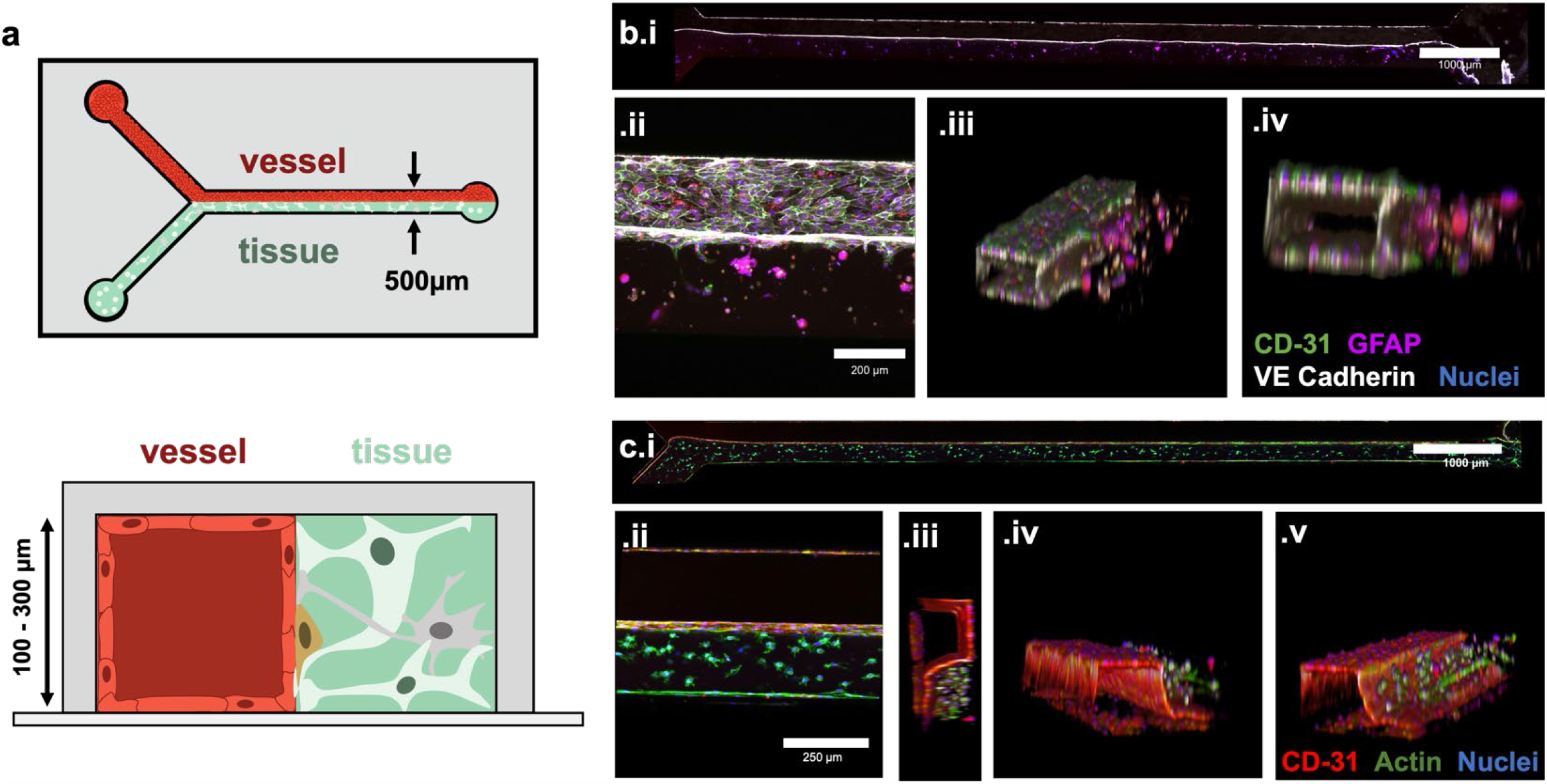
Configuration of blood-tissue interfaces following bioinspiration paradigm in MPS engineering. a) The BTI Chip is a microfluidic blood-tissue interface fabricated within a PDMS microchannel to mimic in vivo microanatomy. A blood vessel endothelium contacts a cell-embedded GelMA-PEG hydrogel. b) Confocal micrographs reveal a blood-brain barrier with hBMVECs alongside an SVGP12-embedded gel. b.i) 3D reconstruction of the SVGP12-embedded gel reveals tissue alignment along the entire 1 cm channel. b.i–iv) Positive staining for endothelial markers CD-31 (green) and VE-Cadherin (white) distinguish hBMVECs from GFAP-positive SVGP12 cells. c) Blood tissue interfaces are customizable. Confocal micrographs reveal an hBMVEC endothelium alongside an HDFn-embedded scaffold. CD-31 staining (red) distinguishes hBMVECs from HDFn. Actin (phalloidin, green) staining shows spreading of HDFn within the tissue scaffold. DAPI (blue) staining indicates cell nuclei.

## 2 RESULTS

### 2.1 Design and Production of BTI Chip

Microchannels were fabricated using soft lithography, while the BTI Chips were generated through a sequential process involving the establishment of tissue compartments using stop-flow lithography, followed by the introduction and culture of endothelial cells. The production flow is shown in **Figure 2**. The Y-shaped microchannel had an inlet channel length of 5 mm and central channel length of 10 mm. The first step consisted of tissue scaffold polymerization along half of the central microchannel by stop-flow lithography. Laminar conditions and the Y shape of the microchannel facilitate a split flow profile in the central channel, so the tissue scaffold precursor and the shaping fluid, *e*.*g*., PBS, each occupy approximately 50% of the central channel. Upon stopping flow, the microchannels were immediately exposed to 5 seconds of UV light. UV light polymerizes the tissue gel scaffold, leaving one side of the central channel occupied by a cell-embedded gel, and the other side filled only with PBS. The flow profiles were monitored and confirmed with an inverted light microscope. Lithium phenyl (2,4,6-trimethylbenzoyl) phosphinate (LAP) was used as the photoinitiator, which has efficient operation across the UV spectrum, up to 405 nm. Accordingly, scaffolds will still polymerize if lower UV-A wavelengths are filtered out in future applications to avoid possible cell damage. After fabrication, endothelial cells were introduced into the open channel, and the devices were placed on their sides in a custom 3D-printed slide holder with the endothelial compartment on top. Devices were held in position for 1 hour to allow hBMVEC to settle on the scaffold wall. Perfusion was introduced after 24 hours to clear the endothelialized lumen and cells were permitted to proliferate for an additional 2 days. This process facilitates the formation of an endothelial cell monolayer. Production and cellularization of the microchannels takes approximately 2-4 days as seen in **Figure 2.c**. These brightfield images illustrate the completed tissue structure and vascular channel. For this report the BTI Chips were fabricated using either HDFn+HUVEC, HDFn+hBMVEC or SVGP12+hBMVEC. HDFn were used for proof-of-principle fabrication, and they were selected due to their rapid proliferation rates, favorable growth behavior in 3D matrices, and compatibility with endothelial cells. Blood-brain barrier (BBB) BTI Chips were manufactured to demonstrate tissue-specific applications. The human astroglia SV40 transformed cell line called SVG p12 (SVGP12) was selected as a BBB-specific cell line due to its conserved properties and its ability to express glial fibrillary acidic protein (GFAP), a hallmark intermediate filament protein in astrocytes, suggesting that key astrocyte characteristics are conserved in this cell line. A detailed description of fabrication and conditions is provided in the *Materials & Methods*.

**Fig. 2.**
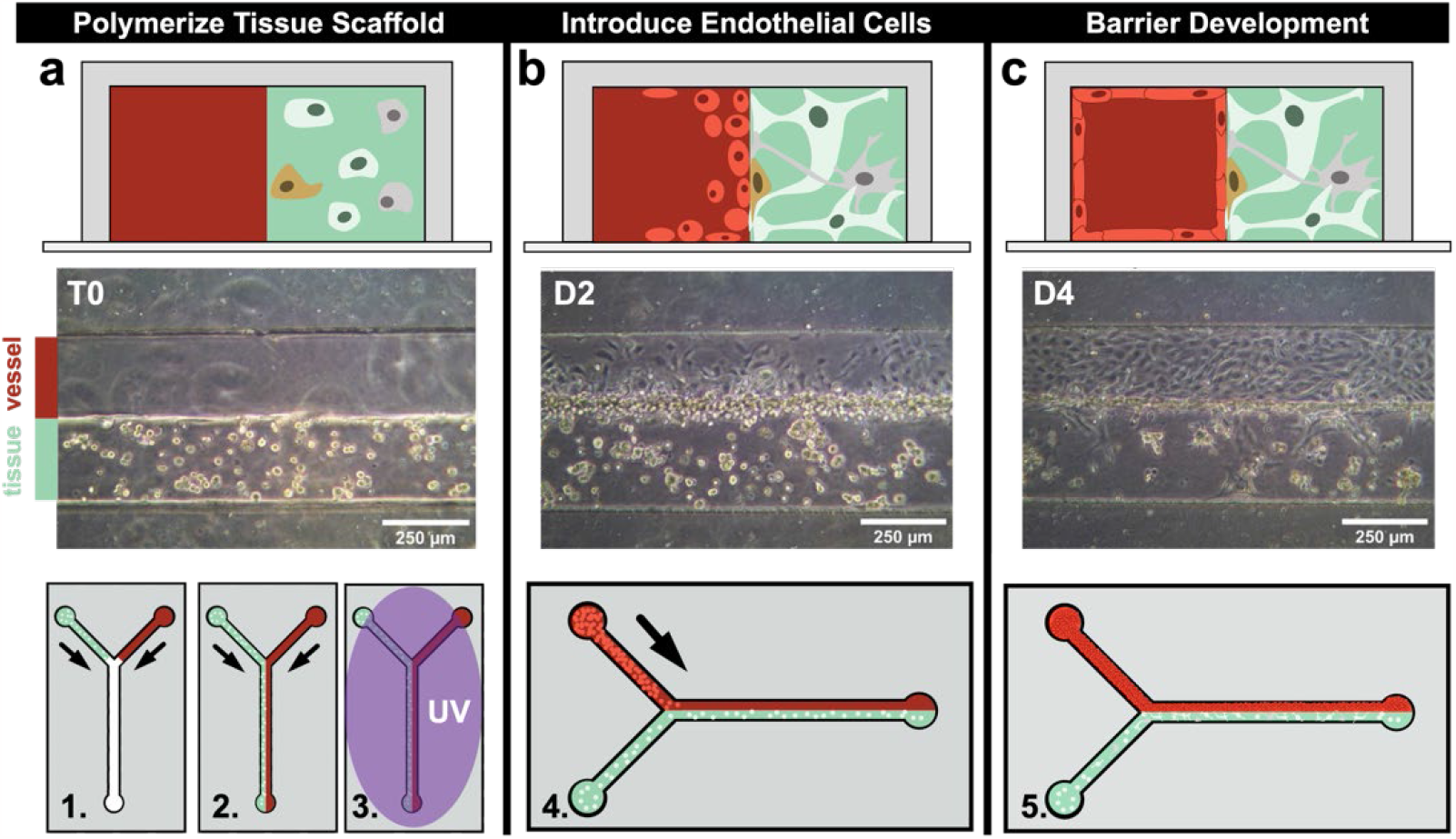
Fabrication of membrane-free Blood-Tissue Interface Chip via stop-flow photopolymerization and endothelialization. a) In steps 1-3, cells suspended in a low wt% mixture of GelMA, PEG-thiol, and photoinitiator are perfused into one of two inlets. The opposing inlet is perfused with 1X PBS such that the cell suspension occupies half of the central channel. The relative split between the blood and 3D tissue compartment can be controlled by the volumetric flow ratios of the input fluids. UV light is applied for 5 seconds immediately after flow is stopped. Microchannels are filled with tissue cell-specific growth medium for 2 days prior to b) adding a high density plug of endothelial cells. Microdevices are temporarily placed on their side to ensure adhesion of endothelial cells to the gel scaffold. c) By day 4, endothelial cells form a complete endothelium and barrier along the tissue scaffold. Brightfield micrographs show the progression of cell culture and endothelium formation.

### 2.2. Verification of BTI Chip Anatomy & Viability

After the construction of the BTI Chip and development of the endothelium in culture, immunohistochemical analysis was conducted to evaluate the organization of the resident endothelial cells and 3D tissue compartment. This confirmed cells from the tissue compartment do not migrate into the blood compartment, and endothelial cells are not mixed into the scaffold during fabrication. Confocal images of the fluorescently labelled MPS are shown in **Figure 1**. Immunohistochemical analysis was utilized to determine the cell location in the MPS and to assess whether tissue cells migrate into the vascular channel or if endothelial cells are driven into the scaffold during fabrication. CD-31, also known as platelet endothelial cell adhesion molecule (PECAM-1), is an endothelial specific protein that makes up a large portion of endothelial intercellular junctions [26]. This protein is selectively labelled in endothelial cells which form a monolayer in the microvascular channel shown in red. F-actin protein is stained green in all cells within the device, which is present in the cytoskeleton of both HDFn and hBMVECs. DAPI staining labels the cell nucleus in blue. Several images were taken along the z-axis of the MPS and max projected to show HDFn proliferation and hBMVEC monolayer formation within the channel as shown in **Figure 1**. hBMVECs adhered on the glass bottom and PDMS top were omitted from the maximum projection image because cell density was increased in these regions, which obscured features in the micrographs. HDFn response to the surrounding matrix is apparent, with dendritic extensions expanding outward towards neighboring cells. hBMVECs occupy the space along the channel and scaffold walls, forming a continuous barrier down the 10 mm central channel in **Figure 1.b.i, 1.c.i**. A clear lumen for media perfusion is shown in the cross-sectional image in **Figure 1.b.iv, 1.c.iii**, though a gap can be seen between the scaffold and the PDMS surface on the top of the microchannel. This gap is caused by the oxygen inhibition of free radical polymerization along the PDMS surface and prevented by oxygen plasma treatment of the microchannel prior to UV polymerization. This gap also contributes to a tapering of the channel wall, which is no longer an issue for shorter channels with oxygen plasma retreatment, as shown in **Figure 3**.

**Fig. 3.**
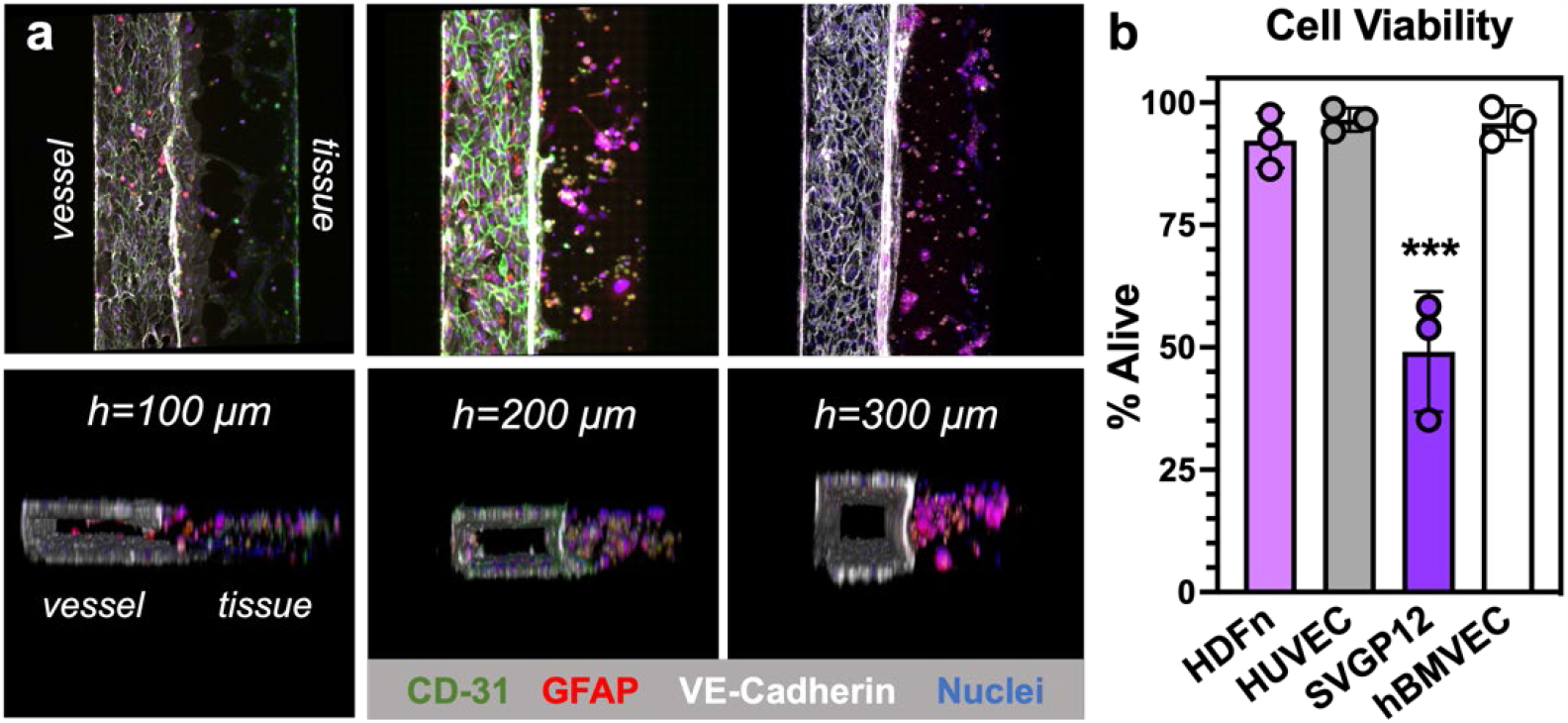
Blood-Interface Chip geometries and post-production analysis of cell viability. a) Confocal micrographs illustrating that 500 µm wide BBB BTI Chips with maintained width of the blood compartment can be fabricated at a variety of microchannel heights (100, 200 and 300 µm). Curvature of the interface is most pronounced above and below heights of 200 µm, which may be attributed to the surface tension and wetting differences among glass PDMS, scaffold materials and media. Illustrative micrographs are BTI Chips consisting of hBMVECs and SVGP12 cells embedded in the 3D tissue scaffold. Staining for CD-31 (green) and VE-Cadherin (white) distinguishes hBMVEC from GFAP-positive (red) SVGP12 cells. All nuclei are DAPI-stained (blue). b) Cell viability was assessed at day 5 of BTI production, one day after addition of endothelial cells. BTI were assembled as HDFn-HUVEC or SVGP12-hBMVEC combinations and maintained under static culture. Data were analyzed via one-way ANOVA (***p<0.001).

The proof-of-concept devices consist of a microchannel with 300 μm height, shown in **Figure 1**. We note that without corona wand treatment prior to addition of gel precursor, there is a tendency for gels to separate from housing material (**Figure 1.c**). Successful attachment to both PDMS and glass is depicted in **Figure 3**. 3D images in **Figure 1.e** show clear definition between the hBMVEC filled microchannel alongside a photopolymerized scaffold, where HDFn can proliferate and form cellular outgrowths in 3D. Differences in scaffold geometry are displayed for BBB MPS with channel heights of 100 μm, 200 μm and 300 μm, shown in **Figure 3**. Channel height of 200 μm shows uniform scaffold formation, as well as a vascular channel width-to-height aspect ratio suitable for shear stress analysis, therefore channels with 200 μm height are used for functional experiments. A complete BTI Chip with SVGP12 glial cells embedded in the tissue scaffold and hBMVECs in the endothelial compartment is shown in **Figure 1.d**. Devices were fabricated with SVGP12 cells mixed into the 3D tissue scaffold for the tissue compartment. Two days after scaffold polymerization, hBMVECs were introduced to the open channel and cultured for two days. The SVGP12 astrocytes exhibited less proliferation, relative to HDFn, as represented in **Figure 1.b and 1.c**, lacking dendritic extensions. Minimal outgrowth of SVGP12 astrocyte projections was observed, but many cells remained spherical, failing to interact with the surrounding matrix. Successful astrocyte interaction would be characterized by end-foot processes covering a large percentage of the microvessel surface. To the best of our knowledge, the recapitulation of interactions between astrocyte end feet and microvasculature in engineered structures has never been reported, suggesting primary and immortalized astrocytes may lose physiological characteristics or struggle to migrate when cultured *in vitro*. Biomimetic astrocyte formation in engineered tissues may be more successful with iPSC technology [27-31]. Nonetheless, astrocytes are shown to conserve glial fibrillary acidic protein (GFAP) production, suggesting key chemokines necessary for BBB maturation are still produced despite morphological differences [32, 33]. Alongside the tissue structure, a complete endothelial monolayer is formed on all channel walls, with junctional proteins present between neighboring cells. The microchannel lumen is conserved, allowing perfusion of media and application of shear stress. The endothelial monolayer and scaffold also remain uniform down the entire 10 mm length of the central channel, shown in **Figure 1.b and 1.c**. There is no gap between the scaffold and PDMS surface after pre-treating channels with corona wand oxygen plasma before fabrication, and the scaffold has a vertical wall without tapering in the z-direction. This uniform geometry is critical for permeability analysis with tracer molecules to ensure limited diffusion and convection of tracer molecules through gaps in the microvascular monolayer. Estimates for the thickness of PBS-GelMA interdiffusion zones that arise prior to UV polymerization are included in Supplementary Information. Flow rate measurements (**Supplementary Information, Figure 1**) at various channel heights were gathered to assist in interdiffusion zone estimation.

Cell viability was assessed to assure UV light exposure and shear forces applied during production of the BTI Chip did not damage cells or induce cell death. UV light exposure has been widely used to crosslink hydrogel scaffolds in tissue engineering with high cytocompatibility with dosages ranging from 138 to 6000 mJ·cm^-2^ [34-39], but UV light (250 nm < λ < 400 nm) raises cytocompatibility concerns due to reactive species generation during exposure, which may cause oxidative DNA damage in embedded cells.[40] The tissue structure is polymerized *in situ* to create the described BTI MPS using a UV light-curing spot lamp that emits energy in the UV-A and visible portion of the spectrum (300-450 nm). The embedded cells in the scaffold are exposed to a UV light intensity of 30 mW·cm^-2^ for 4 seconds during hydrogel polymerization, resulting in a final dosage of 120 mJ·cm^-2^. This dosage is well within the range described in previous studies.

Cell viability in the BTI Chips, with both hBMVEC-SVGP12 and HDFn-HUVEC combinations is shown in **Figure 3.b**. Fluorescence intensity for live and dead stained cells is plotted down the length of the channel in arbitrary units (a.u.), with intensity values averaged along the width of the vessel or tissue structure (**Supplementary Information, Figure S3**). A high live cell fluorescent signal is detected in the vascular channel, consistent along channel length, with low dead cell fluorescent intensity. High viability in the hBMVEC monolayer confirm the applied shear forces do not cause cell death in the vasculature channel. A higher intensity is observed for dead cells along the length of the scaffold, suggesting a lower viability compared to cells in the vascular channel. The sensitivity of SVGP12 to gel solutions was further explored in the Supplementary Information. Results suggest SVGP12 are especially sensitive to the fabrication process, including parameters such as time spent in gel precursor solution prior to UV polymerization.

### 2.3 Regulation of Permeability by hBMVEC Endothelium

Permeation of perfusate from the blood compartment, across the BTI, into the tissue compartment, was assessed by fluorescence microscopy. Fluorescently labelled dextran of different molecular weights was perfused into the blood compartment and the fluorescence intensity realized in the tissue compartment was measured. Experiments were conducted to measure how barrier permeability fluctuates with varying cell types and shear forces. The permeability of a 70kDa Texas Red dextran and 20kDa FITC dextran flowing through a BTI Chip without hBMVECs is compared to the permeability of the tracer molecule in a device with a completely developed endothelium, shown in **Figure 4**. Results demonstrate two-fold greater permeability in devices without hBMVECs compared to SVGP12 and hBMVEC cocultured devices for both 20kDa and 70kDa dextran. These results were expected due to the absence of a barrier layer impeding the convective and diffuse transport of the tracer molecules. Permeability of MPS with SVGP12 cells embedded in the tissue matrix and hBMVECs in the open channel is compared to MPS with an acellular scaffold and hBMVECs in the open channel. No change in permeability was measured between devices with endothelium only versus those co-cultured with SVGP12 for both dextran molecular weights. This may be due to SVGP12 cells lacking key astrocytic protein expression necessary to induce a microvascular response. Previous studies have reported decreased permeability of endothelial cell line monolayers when co-cultured with the SVGP12 cell line, though these studies involved cells adherent in 2D [41, 42]. SVGP12 cells may interact differently when in a 3D culture model, or a higher density of cells may be required to achieve the necessary secreted protein concentration for endothelial response. Representative images obtained during permeability assays are shown in **Figure 4**. Low SVGP12 viability also suggests limited activity and therefore limited contribution to BBB permeability.

**Fig. 4.**
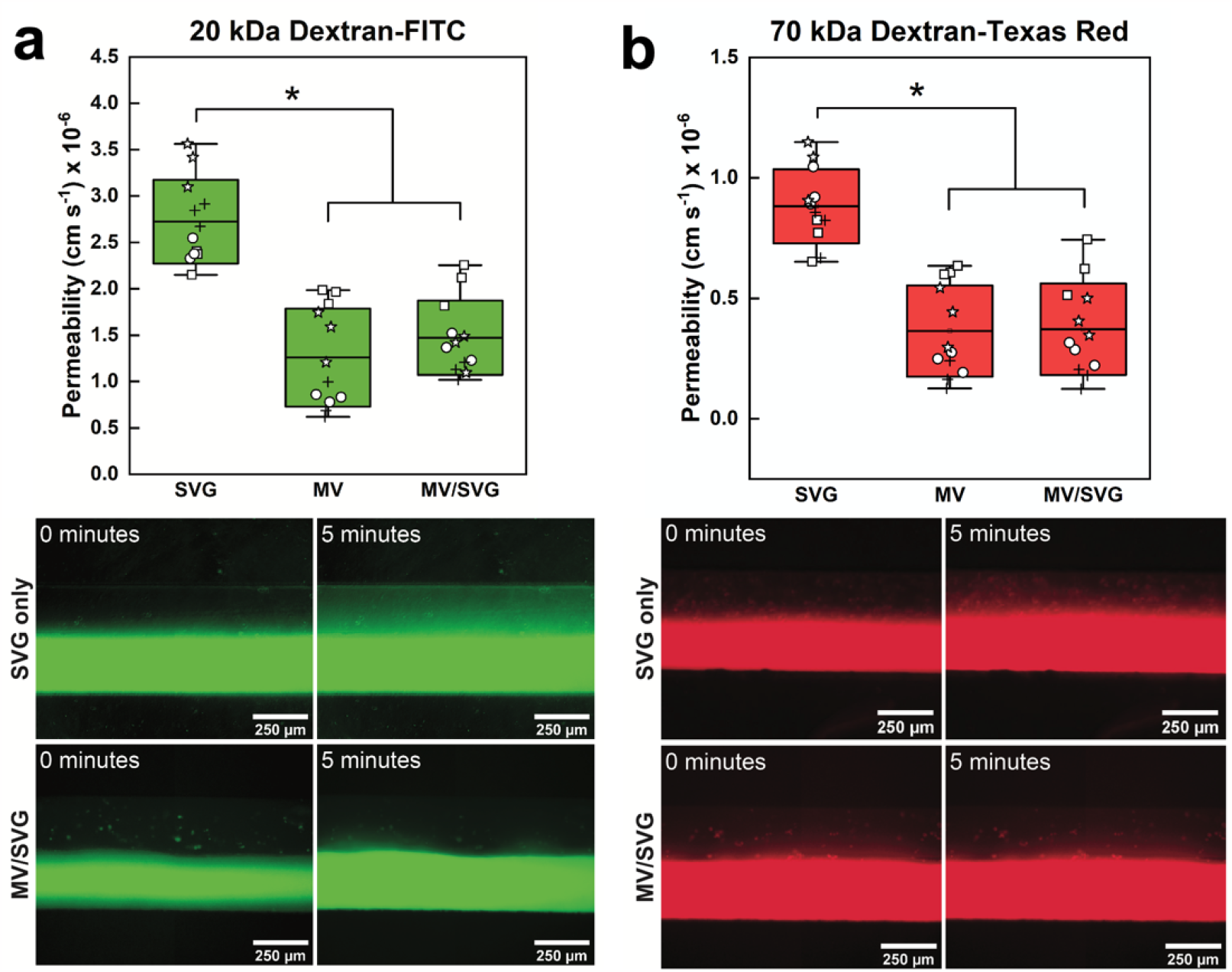
Endothelial barrier was developed and regulated perfusate transport in the Blood-Interface Chip. Fluorescently labelled dextran tracers, (a) 20 kDa and (b) 70 kDa exhibited decreased permeation rate into the 3D tissue compartment when the developed endothelium is present. 20 kDa tracers permeate more rapidly than 70 kDa molecules. Regulation of permeability of the endothelial barrier by the addition of SVGP12 cells into the 3D tissue scaffold was not detected. Each data point represents one of three locations within a single device. 4 devices were used per condition. Data were compared with student’s t-tests, (*p<0.05).

Molecular tracing molecules have been a valuable tool for exploring BBB mechanism of action and response to injury *in vitro* and continue to provide utility in determining physiological relevance of engineered MPS [43-47]. Larger molecules tend to permeate slower across barrier systems, while molecules smaller than 500Da easily diffuse through endothelial layers depending on charge and functionalization. In the BTI Chip, having eliminated the semipermeable membranes, fluorescent tracer molecules can be spatially monitored by standard fluorescence microscopy to calculate permeability over time without sampling that is required in Transwell® or other MPS. In vessels with cylindrical geometries or irregular morphologies in 3D matrices, highly sensitive microscopes must be used to track perfusion in a single plane [48, 49]. Spinning disk confocal microscopy provides the speed and resolution to obtain single-plane images of fluorescent molecules diffusing into a matrix. However, this equipment is not readily available in many research labs, especially considering these assays involve live cells and therefore cannot be transported long distances from culture facilities. Instead, MPS with planar channels and optically transparent windows for imaging, such as the BTI Chip, can be monitored using standard inverted fluorescent microscopes.

### 2.4. Regulation of BBB MPS Permeability by Fluid Shear, Molecular Size, & P-glycoprotein Efflux Pump Activity

While static permeability characterization demonstrated a basal barrier function, the expected operation of the BTI Chip will be under controlled biomimetic perfusion. Accordingly, the permeability was evaluated during extended exposure to perfusion induced shear stress. The permeability of BTI Chips exposed to static and perfusion culture conditions were compared as shown in **Figure 5**. Endothelial cells were exposed to physiological shear stress for 24 hours beginning 1 day after hBMVEC seeding. BBB MPS were exposed to approximately 0 dyn·cm^-2^, 1 dyn·cm^-2^, and 3 dyn·cm^-2^ shear stresses with 0 μL·min^-1^, 5 μL·min^-1^ and 17 μL·min^-1^ flow rates, respectively. Due to media volume and hardware constraints, cells were exposed to shear stress lower than reported physiologically, though similar shear stresses to what are tested herein have been previously reported to decrease permeability in a hBMVEC monolayer treated with astrocyte conditioned media [50]. The permeability of 70 kDa molecular tracers across the BBB MPS does not change with flow, shown in **Figure 5**, as the maximum permeability is achieved with hBMVEC alone due to detection limits and low permeability experienced with large molecules. There is a decrease in permeability of smaller tracer molecules, at 20 kDa molecular weight, across the BBB MPS exposed to shear stresses compared to static BBB MPS, suggesting shear strengthens the junctions between cells and limits permeability, shown in **Figure 5.b**. Shear decreases permeability by 50% compared to BBB MPS without flow, though increasing flow rates to 3 dyn·cm^-2^ does not further decrease permeability. To see a further decrease in permeability of 20 kDa dextran, higher flow rates or longer culture times may be required.

**Fig. 5.**
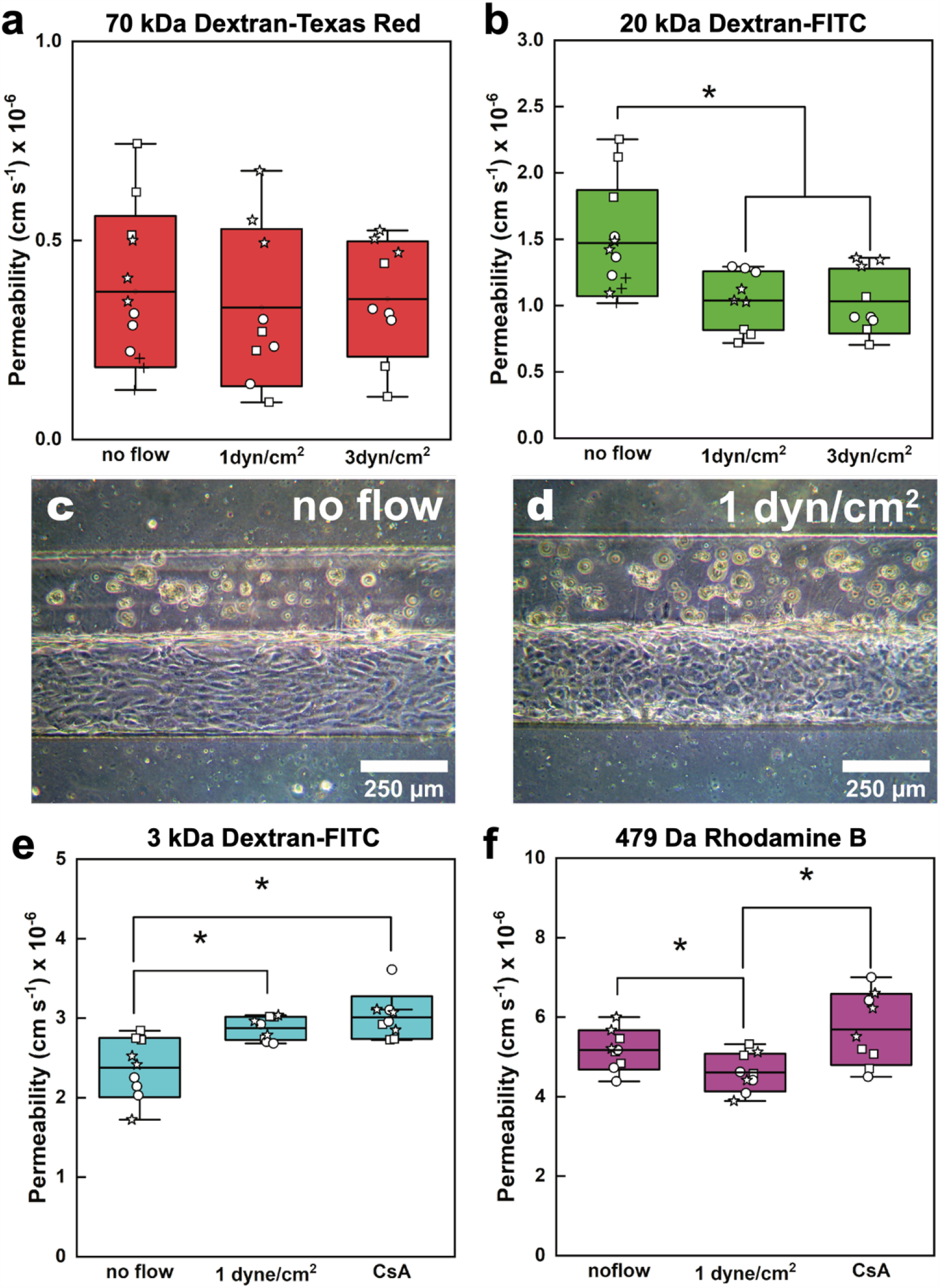
Fluid shear and chemical stimuli regulated endothelial barrier permeability and P-gp efflux pump activity in the Blood-Tissue Interface Chip. a, b) Fluid shear positively regulates barrier integrity and decreases permeability to dextran tracers (20 kDa). c, d) hBMVECs do not elongate in response to shear, confirming brain microvascular endothelial cells do not follow conventional shape-change behavior in response to increased fluid shear. e, f) Inhibition of P-glycoprotein efflux pumps with cyclosporine-A (CsA) increases the net permeability of the endothelial barrier. For rhodamine tracers (479 Da), both shear and efflux pump activity regulated the permeability of the endothelial barrier. Each data point represents one of three locations within a single device. 3–4 devices were used per condition. Data were compared with student’s t-tests, (*p<0.05).

As previously discussed, endothelial cells are known to polarize, elongate, and migrate when exposed to shear stress [51-53]. These key morphological changes highlight the mechanoreceptive capabilities of endothelial cells. Interestingly, microvascular cells from the brain have been reported to show a much different response. Multiple studies have demonstrated how hBMVECs are unlike other types of endothelial cells in that they do not elongate in response to flow [54, 55]. Although no cytoskeletal remodeling is observed, these studies still suggest cells are activated by shear forces. As shown in brightfield images in **Figure 5.c** and **5.d**, hBMVECs in the BBB MPS do not align in the direction of flow with applied shear stress but show alignment without shear. It has previously been reported that multiple cell types, including endothelial cells, align along the length of microchannels, with more alignment observed with decreasing channel width [56, 57]. Similarly, an elongated endothelial morphology is observed along the length of the microchannel in cells cultured statically, as seen in **Figure 5.c**. Upon exposure to flow, this elongation is reduced, and cells appear to have a more squamous morphology. This is due to the unique nature of hBMVEC, where tight junction proteins are critical to prevent intercellular mobility of molecules larger than 500 Da. Shear activated hBMVECs do not elongate to minimize tight junction length between cells, therefore conserving selectivity [54]. hBMVEC reorganization from elongated to squamous morphology is observed, possibly as a mechanism to minimize tight junction length upon shear activation.

The smallest tracer molecule used for permeability analysis was Rhodamine B, a fluorescent tracer dye that has a molecular weight of 429 Da. Molecules smaller than 500 Da can transverse the BBB *in vivo*, though many are subsequently transported back into circulation due to the activity of efflux pumps on the luminal and basal surfaces of endothelial cells. P-glycoprotein (P-gp) is an ATP-dependent efflux pump that limits molecule entry to the brain, presiding on the luminal surface of microvascular cells in the BBB. P-gp is thought to play a role in back-transport of molecules larger than 400 Da [58]. Rhodamine dyes have been widely used to determine functional activity of P-gp, as these molecules are small enough to transverse the barrier to then be actively removed by P-gp efflux pumps [59-62]. *In vitro*, P-gp upregulation has been observed in hBMVECs exposed to shear forces when compared to cells cultured in static conditions [63, 64]. A decrease in permeability of Rhodamine B was seen in BBB MPS exposed to 1 dyn·cm^-2^ shear forces when compared to BBB MPS without shear exposure (**Figure 5.f**) while 3 kDa dextran permeability was the same between flow and no flow conditions (**Figure 5.e**). This may be explained by the upregulation of P-gp on hBMVEC under shear, actively transporting molecules back into the vessel channel as they diffuse into the tissue structure. A subset of the BTI Chips perfused at 1 dyn·cm^-2^ were treated with 10 µM of Cyclosporine-A (CsA) for 1 hour before testing Rhodamine B permeability as seen in **Figure 5**. Permeability of Rhodamine B is shown to increase after BBB MPS incubation with CsA, suggesting efflux pumps that were previously activated by shear stresses are functionally inhibited. CsA is an immunosuppressant drug that has been shown to inhibit P-gp efflux pump activity. CsA is a competitive inhibitor, effectively blocking efflux pump function, leading to the accumulation of Rhodamine B in the scaffold on the basal side of the hBMVEC monolayer. The permeability of Rhodamine B in shear-exposed BBB MPS treated with CsA is the same as permeability seen in BBB MPS cultured without flow, suggesting a physiological response activated by flow is absent after CsA treatments. An alternate explanation for increased Rhodamine B permeability is that CsA treatment physically damaged the BBB. A small increase in permeability of 3 kDa dextran molecules was observed for CsA-treated BBB MPS compared to conditions with and without flow (**Figure 5.e**). An effect is more likely due to barrier damage rather than efflux pump inhibition, as morphological changes were realized at higher dosing of CsA.

### 2.5. Parallel Assembly of Multiple BTI Chips

The described studies run in parallel could provide a wealth of data with limited user requirements, saving time and resources, as well as demonstrating the utility of MPS in settings that require high throughput. Parallel channels of equivalent size will have the same hydrodynamic resistances and pressure drops, ensuring solution will flow simultaneously through all devices. Input pressure must account for the sum of the hydrodynamic resistances across all devices, requiring higher pneumatic pump pressure during fabrication compared to a single channel. Simultaneous fabrication of multiple BTI Chips was achieved as a proof-of-concept for future scale-up. Polymerized scaffolds with embedded HDFn are shown in **Figure 6**. Fibroblasts proliferated in each parallel scaffold, forming dendritic outgrowth into the surrounding matrix, while hBMVECs proliferated to fill the remaining channel in each device. Scaffold polymerization is not uniform in all devices, and resistance differences in each channel after complete MPS fabrication prevented simultaneous perfusion. These limitations could be addressed by using a flood lamp rather than a spot lamp for more uniform UV distribution in large polymerization areas, as well as utilizing stiff, plastic microfluidic channels with PEEK tubing to limit transient fluid motion after stop flow. With this design, there were 12 Y-channels on a 25 mm x 75 mm glass slide. Four of the 12 devices available on the slide were simultaneously fabricated to demonstrate proof of concept (**Figure 6**), though this fabrication can be scaled-up for production.

**Fig. 6.**
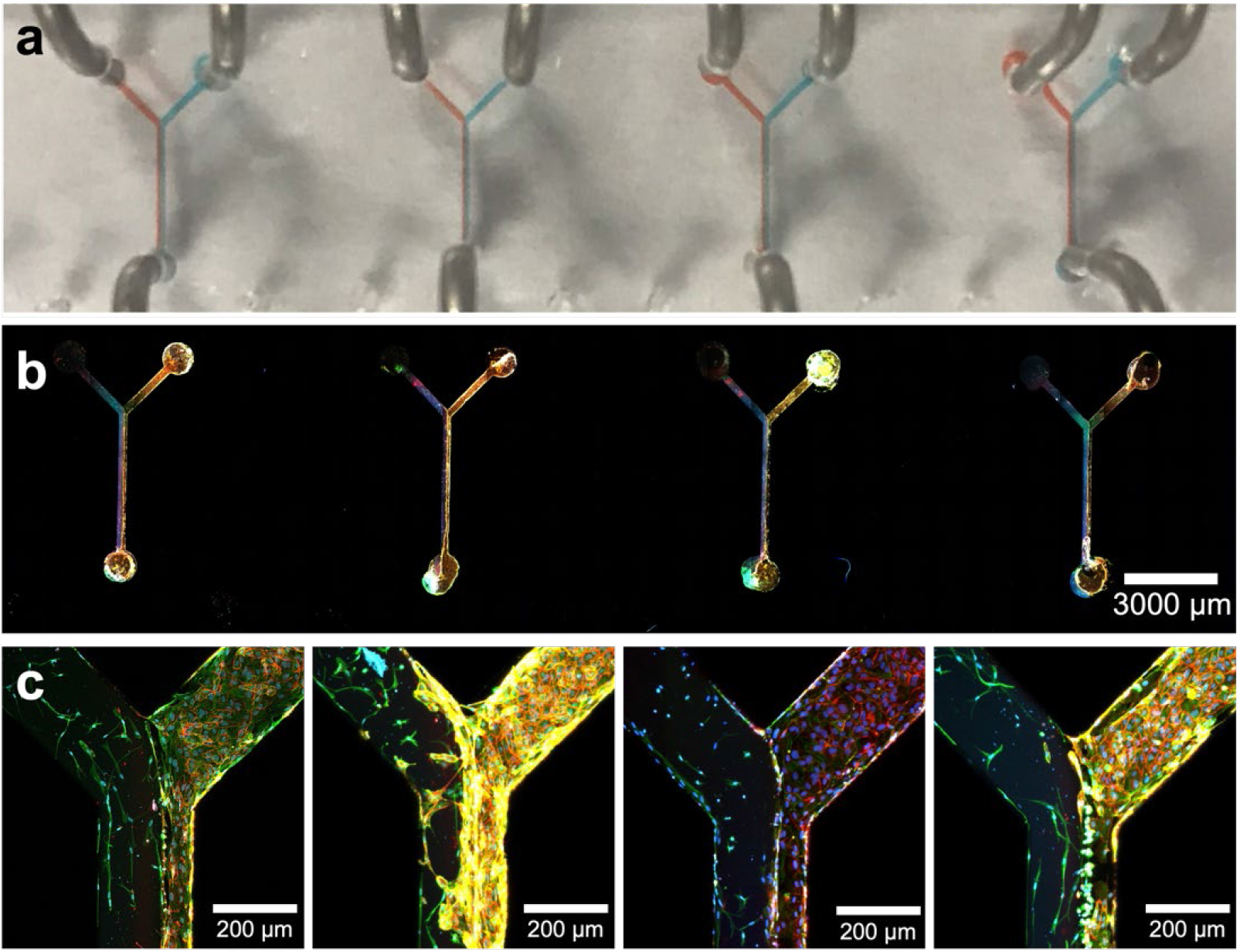
Multiplexed assembly of Blood-Tissue Interface Chip. a) Parallel laminar flow profiles can be established simultaneously in multiple Y channels within the same PDMS housing. b, c) Immunofluorescence micrographs of HDFn-embedded gels and hBMVEC endothelium demonstrate successful fabrication of an array of blood-tissue interfaces. Staining for VE-Cadherin (red) distinguishes hBMVEC from HDFn (phalloidin-stained actin, green; DAPI-stained nuclei, blue). Yellow coloration is a product of red and green fluorescent overlap. All channels are simultaneously constructed and perfusable by simple splitting of the inlet conduits.

## 3. MATERIALS & METHODS

### 3.1. Gelatin methacryloyl (GelMA) Synthesis

GelMA was synthesized as previously reported [65]. Porcine gelatin (Millipore Sigma, Type A, 300 g Bloom) was dissolved in 1X Phosphate Buffered Saline (PBS) (0.1 g·mL^-1^) at 60°C. After the gelatin was dissolved, the solution was cooled to 50°C and methacrylic anhydride (MilliporeSigma) was added dropwise to a ratio of 750 μL methacrylic anhydride per 1 gram of gelatin. The reaction continued for 4 hours at 50°C. The reaction was stopped by adding an equivalent volume of PBS. The solution was dialyzed in deionized water (∼18.2 MΩ·cm^-1^) using regenerated cellulose dialysis tubing with a molecular weight cutoff of 12–14 kDa (SpectrumLabs) at 40°C for 7 days. The dialysate was changed daily. The aqueous GelMA solution was frozen at -80°C overnight and then lyophilized for 5 days. The final product was stored at 4°C until used.

We opted to use GelMA (gelatin methacryloyl) instead of collagen gel for our microdevice fabrication due to its unique advantages. GelMA is a modified form of gelatin, offering greater versatility and mechanical stability compared to collagen. Its incorporation of methacryloyl groups enables the utilization of photopolymerization, transforming GelMA into a hydrogel through UV crosslinking [66, 67]. This feature allows us to precisely control the mechanical properties of the scaffold. Additionally, GelMA provides cell adhesion sites and offers adjustable degradation rates, making it an ideal choice for our specific application. While collagen gel is known for its excellent biocompatibility and bioactivity, GelMA’s tunable and robust properties make it more suitable for our microdevice fabrication needs, ensuring optimal performance and longevity.

### 3.2. Microchannel Fabrication

Microchannels were fabricated by soft lithography [68]. Photolithography masks were designed in AutoCAD (Autodesk) and printed on Mylar® films at 32 kDPI resolution (FineLine Imaging). 100 mm silicon wafers were cleaned with isopropanol (VWR), dried with N_2_, and O_2_ plasma treated (Harrick Plasma) for 2 min prior to coating with SU-8 photoresist (MicroChem, Kayaku) using a spin-coater. SU-8 molds of various heights were created. Baking, exposure, and development times were adjusted according to manufacturer recommendations. After development, molds were hard baked at 150°C for at least 10 min. Polydimethylsiloxane (PDMS) was cast and baked for at least 20 min at 70°C prior to removal from the molds. Inlet and outlet holes were punched in the PDMS devices using an 18-gauge blunt-tip dispensing needle. Dust was cleaned from the PDMS devices using transparent Scotch® tape. PDMS devices and glass slides (25 x 75 mm) were O_2_ plasma treated for 2 min and permanently contact-bonded. The assembled devices were stored without tubing.

### 3.3. Cell Culture

hBMVECs were purchased from Angio-Proteomie (Catalog #cAP-0002) and cultured in microvascular endothelial cell growth media 2 (EGM-2-MV) purchased from Lonza (Catalog #CC-3202). HDFn were purchased from ATCC (Catalog # PCS-201-010) and cultured in DMEM (4.5 g·L^-1^ glucose, L-glutamine, & sodium pyruvate; Corning) supplemented with 10% FBS (Genesee) and 1X penicillin/gentamicin. SVGP12 cells were purchased from ATCC (Catalog #CRL-8621) and cultured in DMEM supplemented with 10% FBS and 1X penicillin/gentamicin. Standard aseptic techniques were used for culture in an incubator with 5% CO_2_ and 100% humidity at 37°C. Cells were cultured in polystyrene culture flasks, media was changed every 2 days, and cells were passaged at approximately 80% confluency. Cells were used to fabricate the BTI Chip up to Passage 6.

### 3.4. Cellularization of Microchannels

#### 3.4.1. In Situ Polymerization of Tissue Compartment

To prepare the BTI Chip, blunt 19G needles (Jensen Global) were removed from plastic Luer Lock fitting by soaking in isopropanol for at least 24 hours. The removed blunt needles were cut to 0.25” using side-cutting pliers and opened using needle-nosed pliers. Blunt needle tips were inserted into channel inlets and outlets, followed by connection to 0.04” ID x 0.07” OD Tygon® tubing. The inner surface of microchannels was treated with a handheld corona treater with antenna attachment for 10 seconds. This was done within 1 hour of scaffold polymerization in the device. The devices were flushed with 70% ethanol and all tubing was clamped with pinch valves. Pinch valves (McMaster-Carr), also called plastic noncontact flow-adjustment valves, are specified for plastic tube outer diameter ranging from 0.09375” to 0.25”. Before use, all tubing was primed with 1X PBS. Leaving one inlet clamped shut, scaffold precursor is introduced to the other inlet, and clamped shut. The scaffold precursor consists of 0.5 wt/wt% lithium phenyl-2,4,6-trimethylbenzoylphosphinate (LAP) (Millipore Sigma) as photoinitiator, 3 wt/wt% GelMA, 1 wt/wt% 8-arm PEG-thiol (Jenkem), and a suspension of the desired cells at 3 million cells·mL^-1^. Once the microchannels and tubing were loaded with 1X PBS and one inlet tube was loaded with precursor scaffold, each inlet tube was connected to one of the PEEK-to-conical tubing adapters, with equivalent lengths of upstream PEEK connected at a T-junction. This T-junction was connected to a flow multiplexor (Elveflow, France), which was connected to a pneumatic pressure regulator (OB-1, Elveflow, France). The pressure regulator was connected to a 50 mL conical fluid reservoir filled with 1X PBS. All tubing was primed prior to making connections. Fluids were perfused under constant pressure (∼12 mbar). When the desired flow profile was achieved, stop flow and UV exposure was initiated for in situ polymerization of the tissue compartment. UV exposure on a conventional inverted microscope was facilitated by placement of an 8 mm liquid lightguide (DYMAX). UV-A (320-390 nm) exposure was measured at ∼40mW·cm^-2^ with an ACCU-CAL 50 radiometer (DYMAX). After the tissue compartment was solidified, tubing was clamped, and the device was removed from the PEEK adaptors. The blood compartment was then perfused with DMEM supplemented with 10% FBS and 1X penicillin/streptomycin using a syringe pump (Harvard Apparatus) at 50 µL·min^-1^ for 30 min then stopped. The media was exchanged each day leading up to the addition of endothelial cells.

#### 3.4.2. Addition of Endothelial Cells and Formation of Endothelium

Two days after fabrication of the tissue compartment and culture, EGM-2-MV media with fibronectin (50 ug·mL^-1^) was introduced to the open channel with a syringe pump and incubated at 37°C for 30 min. hBMVECs (∼ 30 million cells·mL^-1^) were pipetted into Tygon® tubing that was pre-sterilized with 70% ethanol. The hBMVECs were introduced to the open channel at 50 μL·min^-1^. Stop-flow was initiated, and the devices were incubated for 6 hours. The channel was flushed with combined media, 50 v/v% EGM-2-MV, 50 v/v% DMEM, 10 v/v% FBS, and 1X penicillin/gentamicin, leaving adhered hBMVECs. Devices were cultured on their sides. Media was replaced daily or perfused continuously. Device progress was monitored with an inverted light microscope. Complete endothelium formation was achieved after ∼24 hours of culture.

#### 3.4.3. Multiplexing BTI Chips

For the Multiplexed BTI Chip, all fabrication and characterization steps were consistent with one minor alteration, *i*.*e*., all the tissue compartment inlets were connected to one inlet and the same was done with the endothelium compartment inlets. This provided a means to produce the BTI Chips simultaneously.

### 3.5. Cellular Viability Analysis

Viability was determined using a LIVE/DEAD assay (ThermoFisher Scientific). Calcein-AM and ethidium homodimer-1 (EthD-1) were used to indicate live and dead cells, respectively. The BTI Chips were perfused with 4 μM Calcein-AM and 2 μM EthD-1 in media using a syringe pump at flow rates < 30 μL·min^-1^. Devices were incubated for 15 min with cellular stains and imaged using an inverted fluorescence microscope with a mechanical stage (Nikon TE-2000E). The green channel was (λ_ex_ ≈ 480/30 nm and λ_em_ ≈ 535/45 nm) was used to measure Calcein-AM, and the red channel (λ_ex_ ≈ 540/25 nm and λ_em_ ≈ 605/55 nm) was used to measure EthD-1.

### 3.6. Immunohistochemical Analysis of BTI Chip

By perfusion through the BTI Chip, the resident cells were fixed, permeabilized, blocked, and stained *in situ*. The BTI Chips were fixed using 4 v/v% paraformaldehyde in PBS for 15 min at room temperature, permeabilized with 0.5 w/v% Triton-X for 10 min at room temperature, blocked with 2 w/v% bovine serum albumin for 45 min at room temperature, and stained as described below. Between each fixing, permeabilization, and blocking step a washing step with PBS was conducted. BTI Chips containing HDFn and hBMVECs were stained with phalloidin (1:2000 dilution) and a mouse anti-human primary monoclonal antibody for CD-31 (1:200) overnight at 4°C, then washed with PBS, and incubated with a goat anti-mouse polyclonal secondary antibody labelled with AlexaFluor594 (λ_ex_ = 590 nm and λ_em_ = 617 nm) (1:500) overnight at 4°C. BTI Chips containing SVGP12 cells and hBMVECs were stained with mouse anti-human monoclonal antibody for CD31 directly conjugated with fluorescein isothiocyanate (FITC) (1:200), a mouse anti-human monoclonal antibody for GFAP directly conjugated with an AlexaFluor594 fluorophore (1:100), and a rabbit anti-human primary monoclonal antibody for VE-Cadherin (1:200) overnight at 4°C. The next day, these BTI Chips were washed with PBS and stained with a goat anti-rabbit polyclonal secondary antibody conjugated with an AlexaFluor 647 fluorophore (1:500) for 1 hour at room temperature. Finally, for all BTI Chips, cell nuclei were labelled with DAPI nuclear stain and washed with PBS prior to imaging by laser-scanning confocal microscopy (Zeiss LSM 710).

### 3.7. Permeability Analysis in BTI Chip

For all experiments described below, tracer molecules were perfused through the BTI Chip with a syringe pump at 5 μL·min^-1^. As a fluorescent signal appears in the blood compartment, fluorescence micrographs were recorded. Fluorescence micrographs were recorded at multiple locations every 30 sec for 5 min with an inverted fluorescence microscope equipped with a custom mechanical stage and on-stage incubator. Permeability was calculated with the following equation,

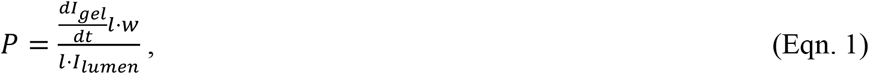

where the change in intensity of the scaffold over the five-minute imaging period, 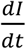, was calculated by taking the slope of the sampled data. *I*_*lumen*_ is the intensity of the lumen at five minutes minus the intensity of the gel at t = 0. The length and width of the scaffold viewing area are *l* and *w*, respectively. For all permeability experiments, 3 locations were imaged per device. 3–4 devices were used for each experimental condition. Images were analyzed using ImageJ [69]. Outcomes were compared with type 2, 2-tail student t-tests. Significance was determined as p<0.05.

#### 3.7.1. Permeability With and Without Endothelium

FITC-labelled dextran (20 kDa; λ_ex_ = 490 nm and λ_em_ = 525 nm) and Texas Red-labelled dextran (70 kDa; λ_ex_ = 596 nm and λ_em_ = 615 nm) in EMG-2 MV media was perfused at 12.5 μg·mL^-1^.

#### 3.7.2. Permeability With and Without Fluid Shear

BTI Chips were exposed to no shear or 1 dyn·cm^-2^. To achieve the desired shear, all perfusion was initiated at a flow rate of 2 μL·min^-1^ (0.4 dyn·cm^-2^), and the flow rate was gradually increased in a stepwise function at ∼2 μL·min^-1^ every 30 min. Applied shear stress of 1 dyn·cm^-2^ was achieved after 1.5 hours, while applied shear stress of 3 dyn·cm^-2^ took 3 hours. Cells were exposed to shear stress for 24 hours. After applying shear stress, permeability was assessed using tracer molecules, as previously described. The fluid shear stress on the endothelium was calculated with the following equation,

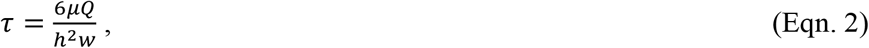

where, *μ* [Pa·s^-1^] is the dynamic viscosity, *Q* [μL·s^-1^] is the volumetric flow rate, *h* [cm] is the channel height and *w* [cm] is the channel width.

#### 3.7.3. Small Molecule Permeability & P-glycoprotein Efflux Pump Activity

BBB MPS were exposed to no shear or 1dyn·cm^-2^. A subset of the MPS perfused at 1dyn·cm^-2^ were treated with 10µM of Cyclosporine-A (CsA) to challenge the P-gp efflux pump for 1 hour before testing Rhodamine B permeability (**Figure 5**). FITC-labelled dextran (3 kDa; λ_ex_ = 490 nm and λ_em_ = 525 nm) and Rhodamine B (429 Da; λ_ex_ = 553 nm and λ_em_ = 627 nm) were introduced to the BTI Chip at 12.5 μg·mL^-1^ in media and monitored every 15 seconds for 5 min, due to the rapid transport of the small molecule, Rhodamine B.

## 4. DISCUSSION

We have demonstrated the fabrication of a novel, microfluidic blood-tissue interface MPS. With our vision for the device, we anticipate achieving a throughput ranging from dozens to hundreds, and through the implementation of an optimized production process, we theorize that production could scale up to an industrial level comparable to Mimetas. Through the strategic use of laminar flow and photopolymerization, we have managed to embed cells in a tissue scaffold that comes into direct contact with a perfusable endothelium, without the need for any synthetic barriers. By incorporating both brain microvascular endothelial cells and model glial cells, we have successfully demonstrated that the BTI Chip can serve as a blood-brain barrier. Our experiments involving the perfusion and tracking of various sizes of fluorescent dextran molecules have shed light on the critical role played by the vascular endothelium and fluid shear response in regulating permeability across the interface. Furthermore, we have proven that permeability across the interface can be modulated via competitive inhibition of efflux pumps using cyclosporine-A. We have also managed to fabricate a parallel assembly of BTI Chips, which holds immense promise for applications in fundamental biological studies and drug development. With MPS already being used for drug toxicology testing on a large scale [3, 4, 11, 12, 23, 70], the automated handling and high number of replicates offered by the BTI Chip’s parallelization of fabrication via microfluidics and photopolymerization are crucial in establishing the utility of microdevices as research and development tools, especially in comparison to animal models and testing.

The geometric ratio of blood and tissue compartments investigated within the BTI Chip was 1:1. Prior work suggests that various width ratios could be achieved by varying relative inlet flow rates [71]. While this approach could provide for a novel method for changing fluid shear in endothelial lumen, the degree of compartment interaction or recapitulation of barrier function is more dependent on the barrier area, which is tuned primarily by height. More exciting derivatives of the BTI Chip would be comparison or combination of tissue cells, or inclusion of a third compartment.

Although UV photoactivatable gels played an important role in the development of our MPS, it is important to recognize the limitations associated with this technique, especially if working with sensitive cell types such as stem cells and some primary cells. Photoinitiators and UV exposure can cause cellular damage, significantly impacting cell viability and functionality. Already, we have seen a relatively high sensitivity of SVGP12 to gel and UV conditions, compared to HDFn. Furthermore, prolonged exposure to UV radiation can result in phototoxicity and disrupt the overall cell culture environment [72]. Therefore, careful consideration of UV exposure parameters, such as time and intensity, is crucial to minimize adverse effects. Alternative crosslinking methods should be explored to address these concerns and preserve cell functionality in future iterations.

An obstacle for many researchers who use cell suspensions of hydrogel precursor solution is the tendency of cells to settle due to gravity. This requires regular mixing of cell-gel suspensions throughout a given fabrication period. In our case, this means mixing in between polymerization of individual BTI Chip gels. Ultimately, cell settling could place a limitation on the number of simultaneously polymerized gels that could be prepared in a multiplexed BTI Chip if gel loading is not done in a single step. More BTI in a single multiplexed chip would require greater gel volume and more loading time. It might therefore be more practical to pursue higher throughput fabrication with automated fluid handling systems, rather than or in addition to multiplexing microfluidic designs. The organ chip company Emulate, Inc. has taken this approach by developing their own ‘Zoë’ chip-handling systems that regulate temperature and CO_2_ as well as media perfusion and stretch of actuator channels. Future experiments aimed at exploring the limits of multiplexing our BTI design are warranted. It is also worth mentioning that MPS research could benefit from experiments exploring the limits of informative capacity for particular MPS dimensions. For example, it would be valuable to answer what the fundamental differences are between deriving information from 12 MPS with a particular channel length “*l*” versus 6 MPS with channel lengths equivalent to “2*l*.” An interdisciplinary approach is warranted to balancing requirements, such as cell counts needed to achieve statistical relevance in a particular assay, with engineering limitations, including incorporating microchannel dimensions that allow realization of uniform fluid shear stress, and pharmaceutical considerations, such as using the minimum necessary amount of material and cells required per device to inform drug performance.

## 5 ACKNOWLEDGEMENTS

A.T.Y was supported by a Pre-Doctoral Training Program in Integrative Vascular Biology at the University of North Carolina at Chapel Hill (NIH Grant 2T32HL069768-16). H.D was supported by the Comparative Medicine Institute and a Pre-Doctoral Training Program at NC State University (NIH Grant T32-GM133393). G.R was supported by the Chemistry of Life Pre-Doctoral Training Grant at NC State University (NIH Grant 5T32GM141887-03). This research was supported by the National Science Foundation (CCSS-1846911, EBMS-2211404, and BMAT-1847488), the American Heart Association (22TPA969368), and the NC State Chancellor’s Innovation Fund. This work was performed in part at the Chapel Hill Analytical and Nanofabrication Laboratory, CHANL, a member of the North Carolina Research Triangle Nanotechnology Network, RTNN, which is supported by the National Science Foundation (ECCS-1542015), as part of the National Nanotechnology Coordinated Infrastructure (NNCI).

