## Supplemental Information for "Simple Design for Membrane-Free Microphysiological Systems to Model the Blood-Tissue Barriers"

When perfusing separate fluids into the inlets of a Y or T junction, some diffusion is expected between fluids. Multiple simulations and experiments have demonstrated that more mixing is achieved at greater channel lengths and with greater aspect ratios of height and width [1, 2]. At low height-width aspect ratios and relatively high Peclet numbers achieved with higher velocities, the amount fluid-fluid interdiffusion is minimal. For a particular diffusion coefficient, D, we can approximate the time to establish mixing in a channel of width *w* with:

$$t = w^2/4D$$

Accordingly, if we know the perfusion velocity, we can calculate the width of mixing at the time it takes to reach our maximal channel length (1 cm). At a supplied pressure of 15mbar, if D is estimated as $1\times10^{-10}$ for the diffusion of gelatin methacryloyl (GelMA) in water. Using

$$Q=vA,$$
$$v=d/t,$$

and our measured flow rate values for various channel heights (100 µm, 200 µm, and 300 µm) from Figure S1, we can calculate the velocities as 4.29 cm/s, 4.24 cm/s, and 3.7 cm/s, and the corresponding maximal interdiffusion mixing widths, at the full channel length of 1 cm, as 9.7 µm, 9.7 µm, and 10.4 µm, with width constant at 500 µm. It is also worth noting that conditions of interdiffusion change when flow is stopped and UV applied. When stop-flow polymerization occurs, the gel flow rate slows at a greater rate than the counterflow fluid, which could lead to an amount of interdiffusion that differs from predictions. At predicted interdiffusion widths, the bulk characteristics of the hydrogel remain largely consistent. While embedded cells should experience uniform substrate mechanics, endothelial cells attached to the surface of the hydrogel would experience a gradient of stiffness from junction to outlet. The impact of this mechanical gradient is not evaluated herein, but Y and T junction microchannels present themselves as a potential platform for investigating cellular responses to substrate with mechanical characteristic gradients. Previous work indicates cells such as fibroblasts do respond to gradients with durotaxis, the migratory preference for stiffer surfaces [3]. Substrate stiffness has also been implicated in cell monolayer permeability [4], suggesting it could be possible to generate an endothelial barrier with a gradient of permeability, provided the correct mixing conditions of gel precursor solution and counterflow fluid prior to gel polymerization in a Y channel. Fluorescent intensity values for BBB permeability calculations were collected along three locations within each device to capture average permeability.

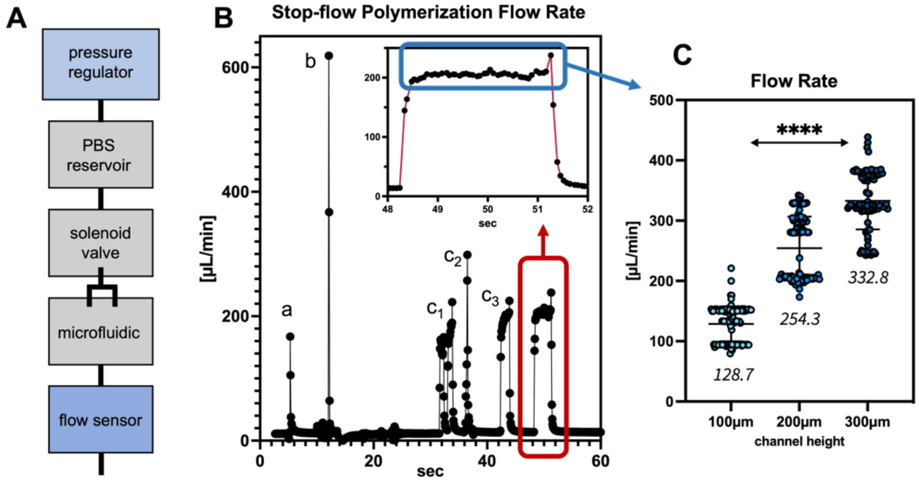


**Figure S1. Regulating and tracking flow during in situ polymerization of tissue compartment gels within Y channel microfluidics.** *Flow values were tracked with an SLI-1000 Sensirion flow sensor connected downstream of a Y channel microfluidic (A). B) Spikes in flow rate values at 'a' and 'b' correspond to unclamping tubing clamps downstream of a sensor and upstream of a microfluidic chip, respectively. Rise and fall in flow rate at '$c_{1-3}$' are from opening and closing a solenoid valve (A) that connects the microfluidic to pressure (15 mbar applied). A sequence of three pressure applications is a standard approach to confirming split flow profile prior to applying UV. The final rise and fall in flow rate correspond to a final opening and closure of the solenoid; UV is applied immediately upon closing the solenoid. Flow rate values from the final pressure spike (B) are averaged for a group of devices by height (C). Flow rate increases with microchannel height due to the decrease in microchannel resistance. Data in C is presented as Sensirion Sensor values collected at 16-bit resolution and 115200 baud rate. 3-4 devices were used for each height; data was compared via one-way ANOVA (****p<0.0001).*

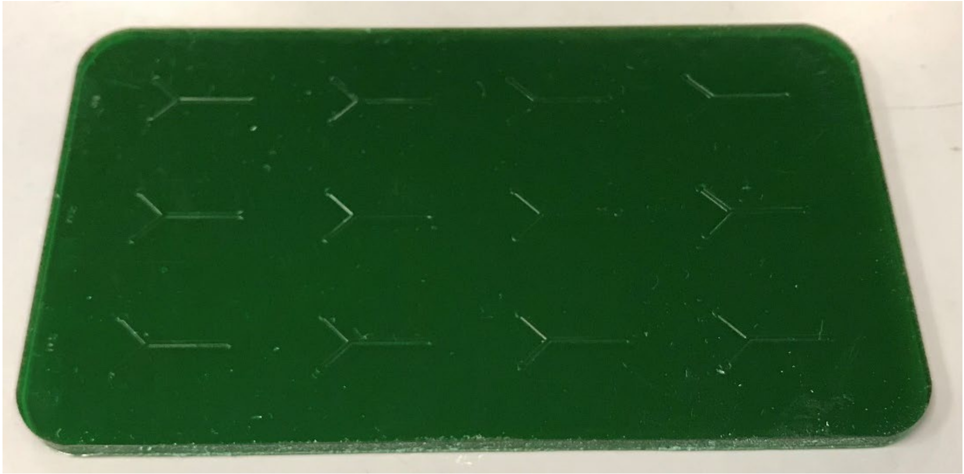

**Figure S2. 3D-printed microfluidic Y-channel mold for casting PDMS microchannels.** *This mold was designed in AutoCAD and printed with a CADworks3D ProFluidics 285D microfluidics printer at a 50 µm z-slicing resolution.*

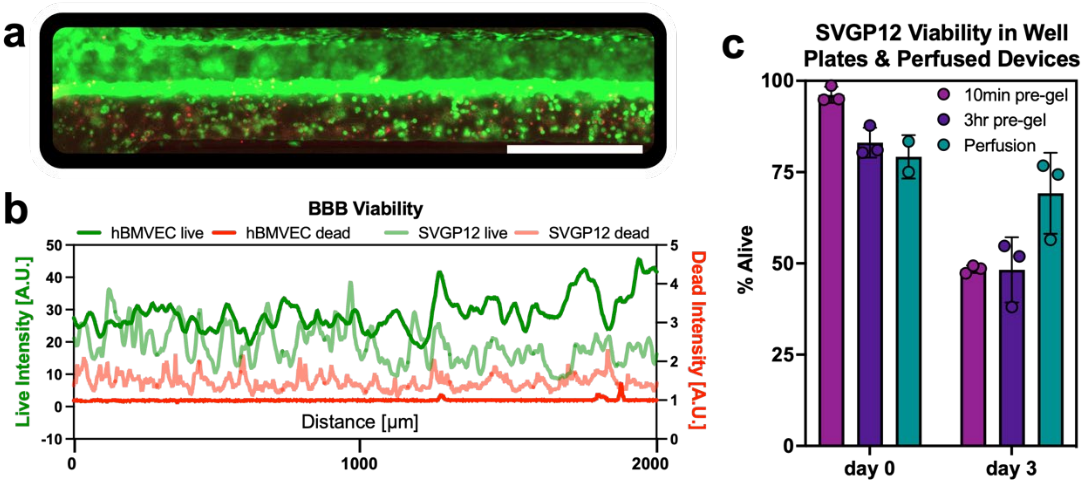


**Figure S3. SVGP12 and hBMVEC Viability.** *A fluorescent micrograph (A, scale bar: 500µm) and the corresponding relative fluorescent intensities of live and dead SVGP12 and hBMVEC (B) are presented for a BBB BTI Chip. Additional viability tests were performed with SVGP12 in well plates and in microchannels (C). Instead of static culture, media was perfused at ~16.5 µL·min*$^{-1}$. *n=2-3 for each condition.*

We further probed SVGP12 viability with LIVE/DEAD assays (ThermoFisher Scientific), where calcein-AM and ethidium homodimer-1 (EthD-1) were used to indicate live and dead cells, respectively. We tested the impact of time spent in pre-gel suspension, prior to UV polymerization. We also compared static and dynamic culture, i.e., continuously replenishing growth media via perfusion of microchannels. Cells were resuspended at ~4 million·mL$^{-1}$ in gel precursor solution. To test the impact of time spent in pre-gel, cells were incubated in liquid GelMA solution at 37 ºC for 10 min or 3 hr prior to polymerization. 50 µL of cell-gel mix was added to each well of a flat-bottom 96-well plate (VWR). Plates were exposed to ~25 mW·cm$^{-2}$ for 5 s (~125 mJ·cm$^{-2}$). Within 2 hr of polymerization or 3 days after polymerization, cells were dyed with LIVE/DEAD dye for 15 min prior to rinsing with 1X PBS and imaging of 150 µm z-stacks on a Leica DMI-8 Stellaris 5 confocal microscope at 1.5 µm z-slicing. Live and dead cells were quantified via ImageJ (NIH) by creating maximum projections, separating channels, thresholding, binarizing, and counting relative green and red particles. This process was repeated for SVGP12 polymerized in 200 µm microchannels; some devices were dyed with LIVE/DEAD dye within 2 hr of polymerization and the remaining devices were dyed after 3 days of microchannel perfusion with a peristaltic pump (Ismatec). DMEM at 10% FBS and 1X penicillin/streptomycin was perfused at ~16.5µL·min$^{-1}$. For "day 0" devices, LIVE/DEAD dye introduced at 50 µL·min$^{-1}$ and given 15 min under static conditions at room temperature prior to rinsing with 1X PBS at 50 µL·min$^{-1}$. For "day 3" devices, cells were dyed with DAPI and annexinV-TRITC to distinguish total and dead cell counts, respectively. The gels were imaged as z-stacks and images processed as described. Results are presented in Figure S3. Compared to devices maintained under static culture, devices with continuous replacement of media achieved a higher viability (69% vs 49%). Each of these viabilities are relatively low but might be improved via alternative photopolymerization mechanisms or alternative tissue gel materials.
